## Supplemental Figures S1-S7 for "Methylome evolution through lineage-dependent selection in the gastric pathogen *Helicobacter pylori*"

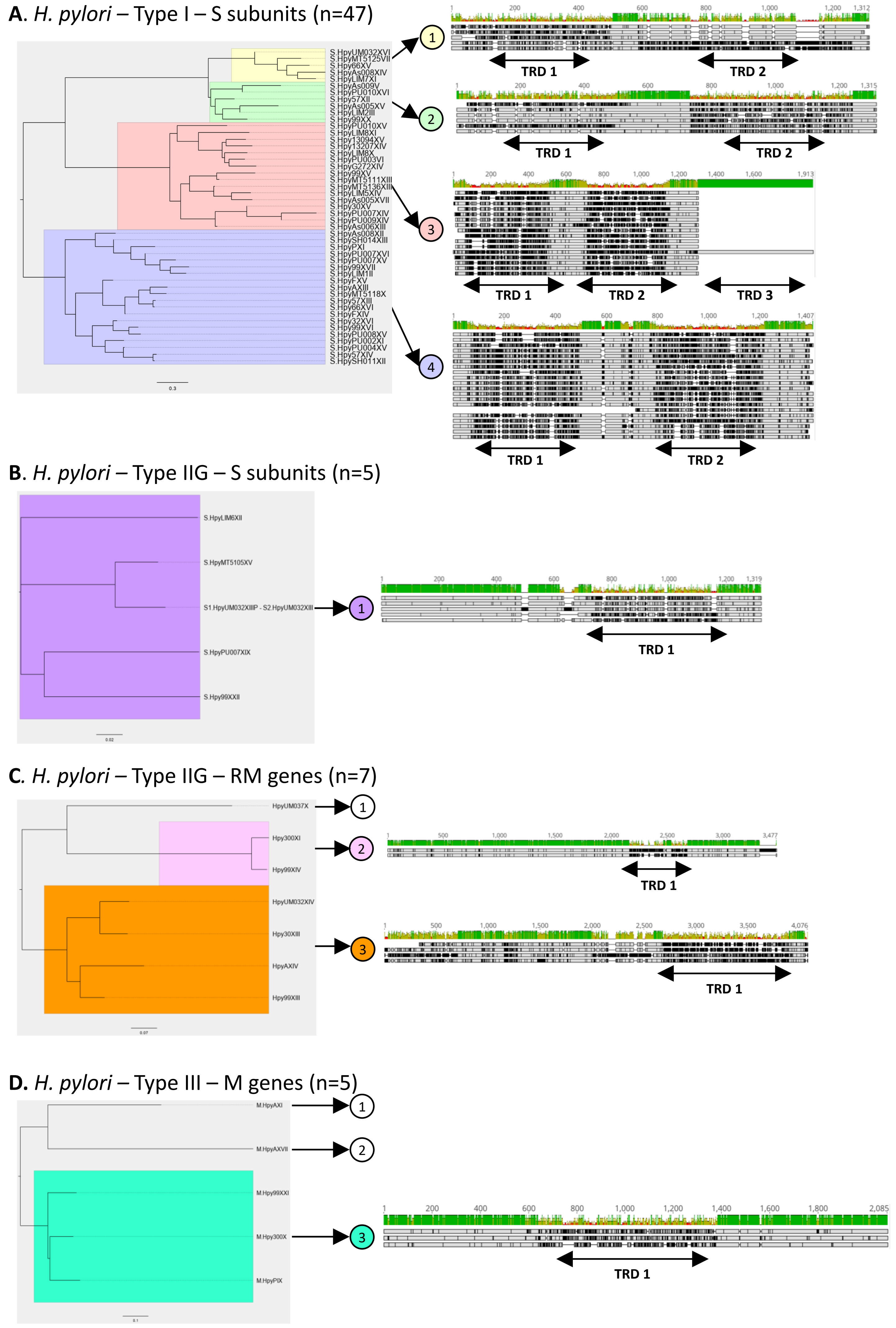


Figure S1 – Phylogeny of type I, IIG, and III RM systems in *H. pylori*.

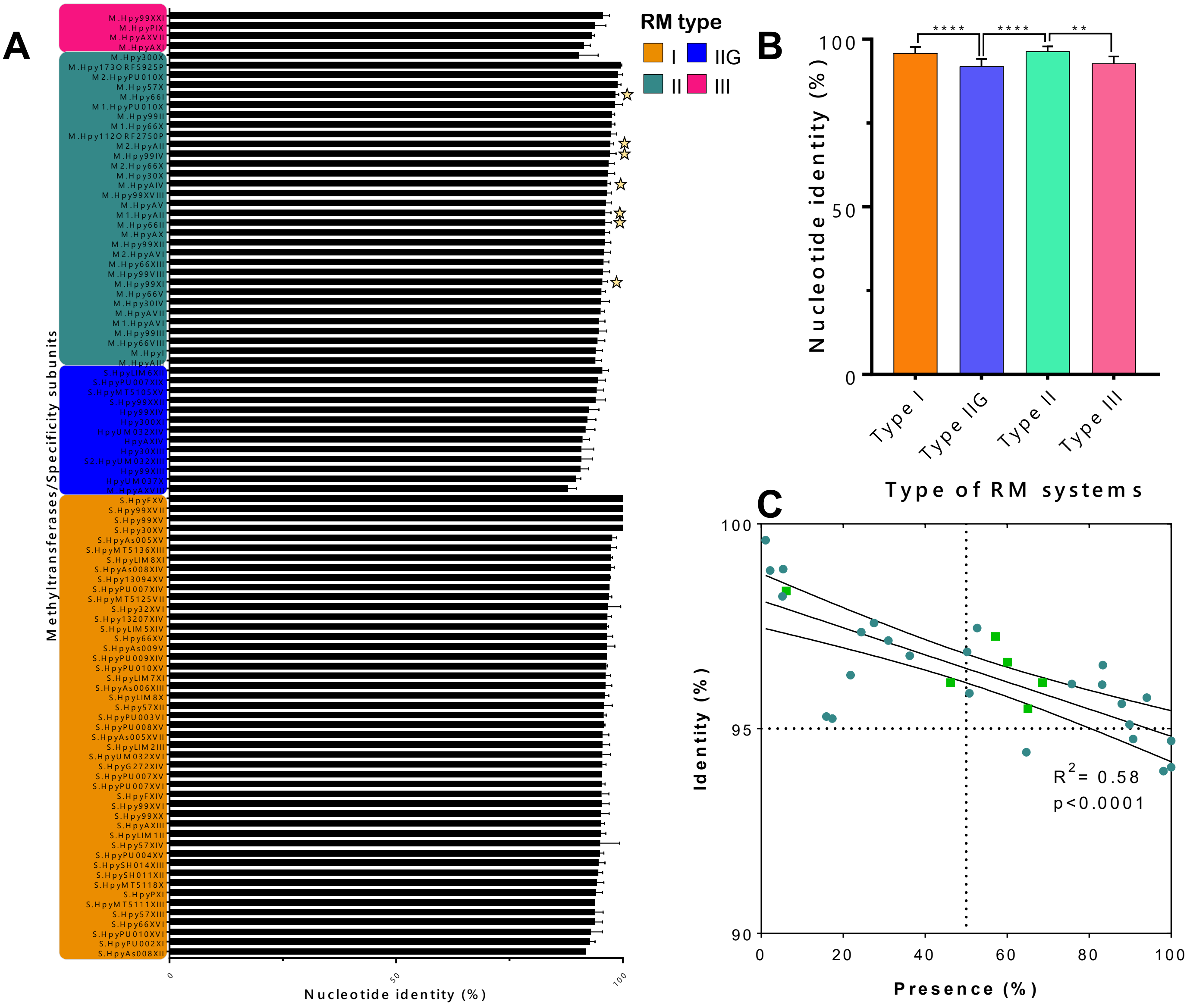


Figure S2 – Sequence conservation of methyltransferases in *H. pylori*.

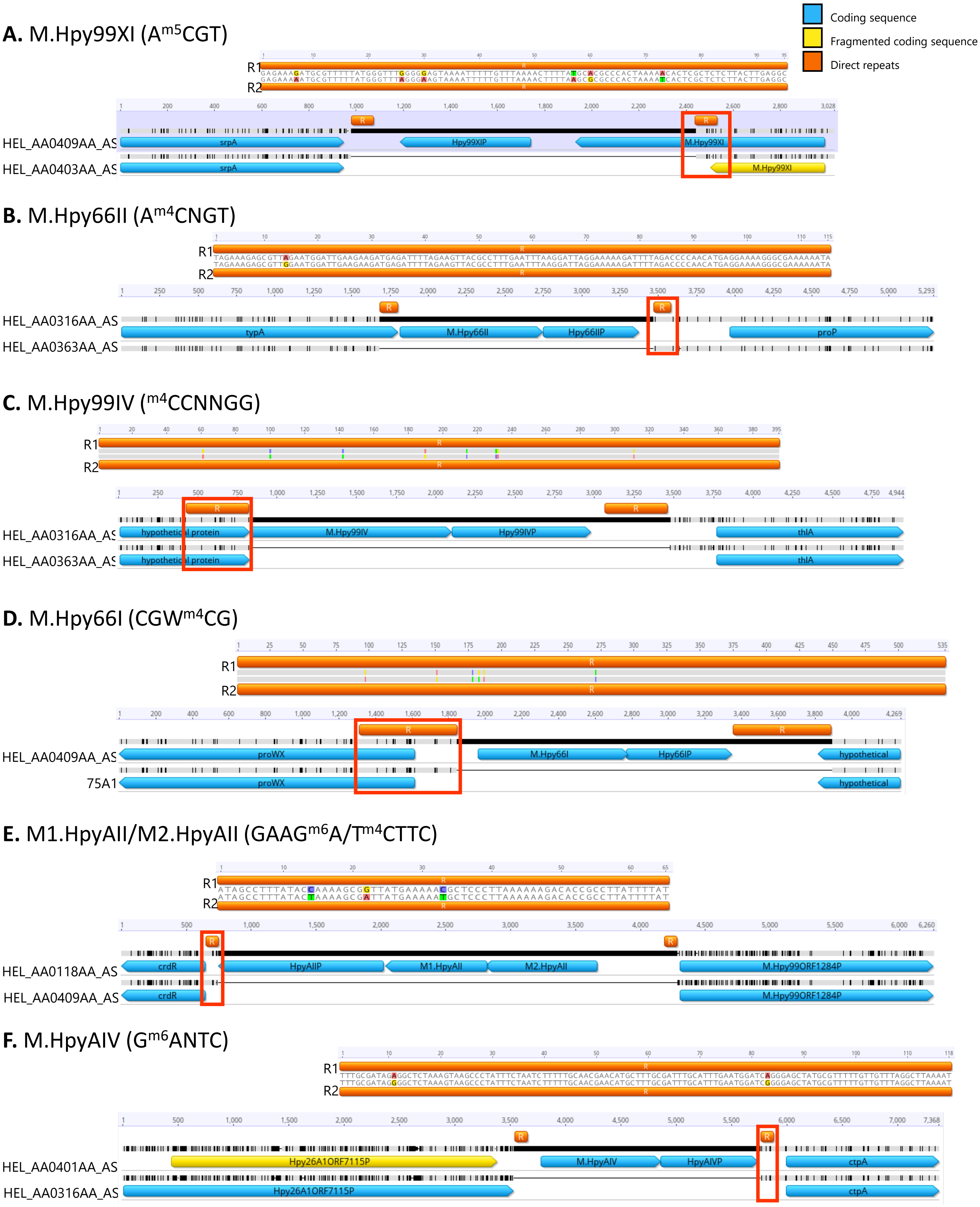


Figure S3 – Direct repeats flanking six different type II RM systems.

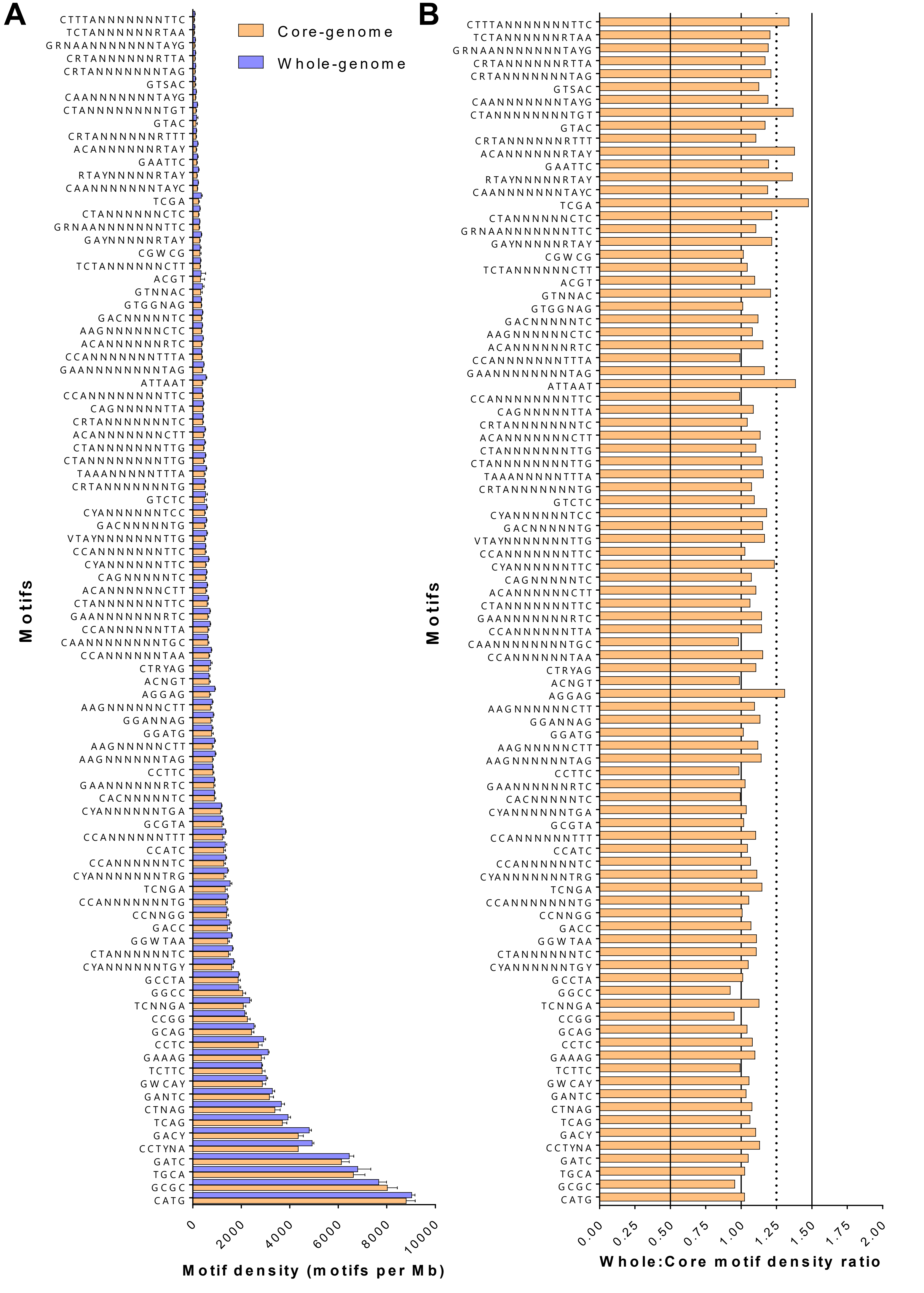


Figure S4 – Comparison of target motif density in whole genome and core genes.

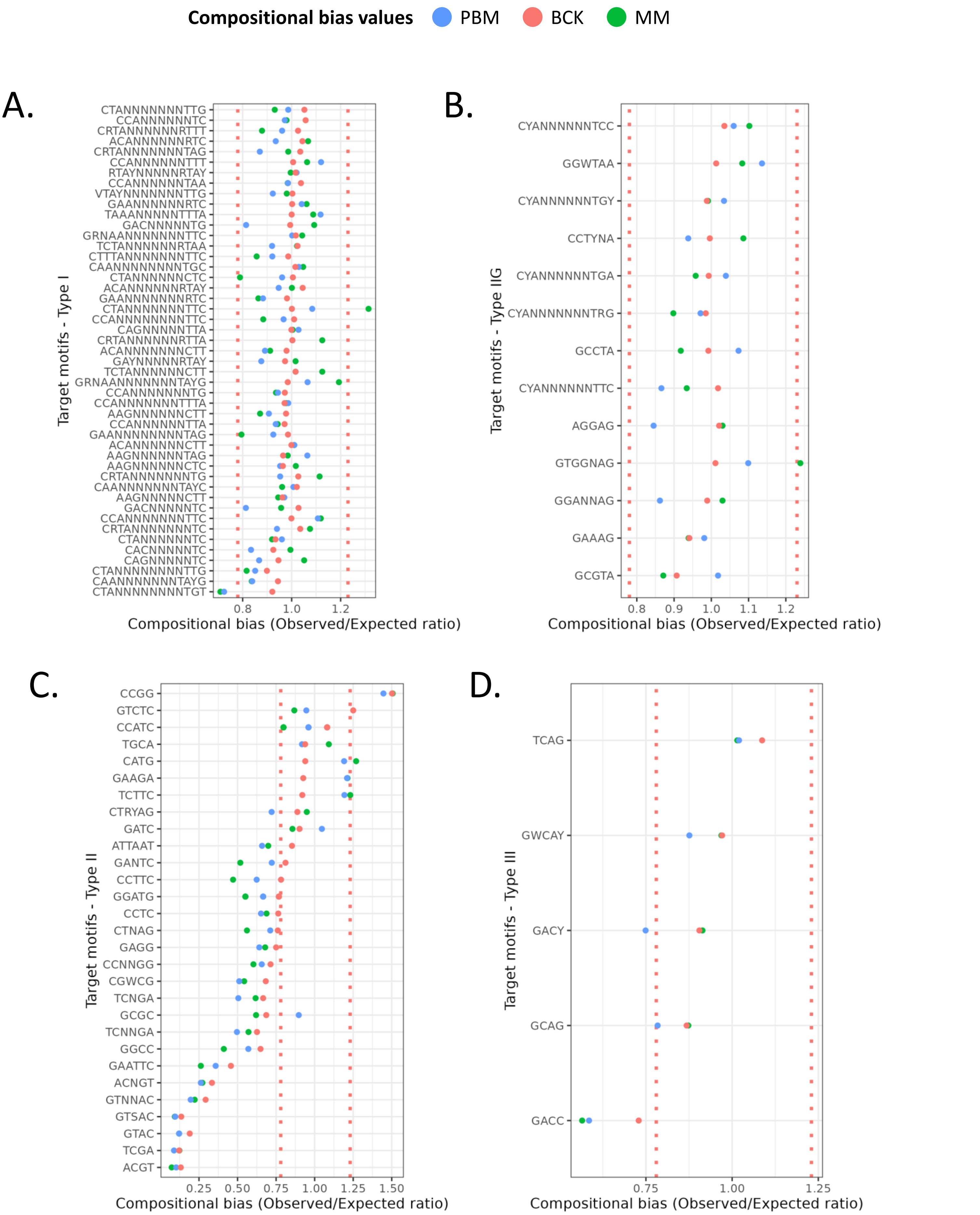


Figure S5 – Compositional bias calculation for target motifs in *H. pylori* using three different methods.

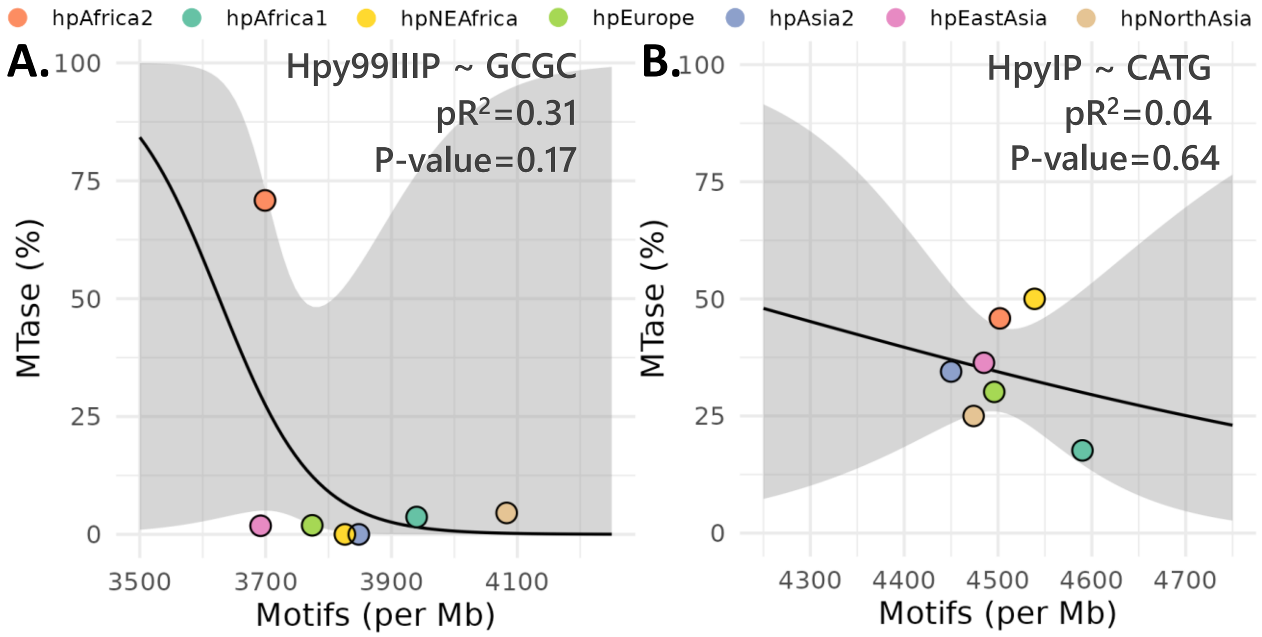


Figure S6 – Interaction between endonuclease frequency and motif density in two type II RM systems.

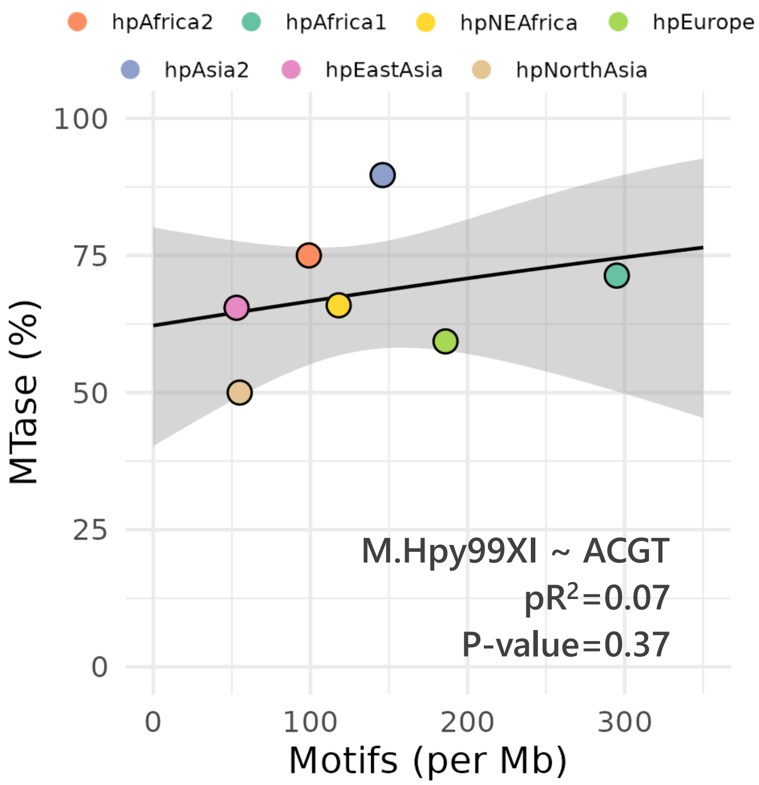


Figure S7 – Interaction between methyltransferase frequency and motif density for the Hpy99XI RM system.
